# Opposing Motor Memories in the Direct and Indirect Pathways of the Basal Ganglia

**DOI:** 10.1101/2024.02.26.582159

**Authors:** Kailong Wen, Zhuoyue Shi, Peijia Yu, Lillian Mo, Shivang Sullere, Victor Yang, Nate Westneat, Jeff A Beeler, Daniel S McGehee, Brent Doiron, Xiaoxi Zhuang

## Abstract

Loss of dopamine neurons causes motor deterioration in Parkinson’s disease patients. We have previously reported that in addition to acute motor impairment, the impaired motor behavior is encoded into long-term memory in an experience-dependent and task-specific manner, a phenomenon we refer to as aberrant inhibitory motor learning. Although normal motor learning and aberrant inhibitory learning oppose each other and this is manifested in apparent motor performance, in the present study, we found that normal motor memory acquired prior to aberrant inhibitory learning remains preserved in the brain, suggesting the existence of independent storage. To investigate the neuronal circuits underlying these two opposing memories, we took advantage of the RNA-binding protein YTHDF1, an m^6^A RNA methylation reader involved in the regulation of protein synthesis and learning/memory. Conditional deletion of *Ythdf1* in either D1 or D2 receptor-expressing neurons revealed that normal motor memory is stored in the D1 (direct) pathway of the basal ganglia, while inhibitory memory is stored in the D2 (indirect) pathway. Furthermore, fiber photometry recordings of GCaMP signals from striatal D1 (dSPN) and D2 (iSPN) receptor-expressing neurons support the preservation of normal memory in the direct pathway after aberrant inhibitory learning, with activities of dSPN predictive of motor performance. Finally, a computational model based on activities of motor cortical neurons, dSPN and iSPN neurons, and their interactions through the basal ganglia loops supports the above observations. These findings have important implications for novel approaches in treating Parkinson’s disease by reactivating preserved normal memory, and in treating hyperkinetic movement disorders such as chorea or tics by erasing aberrant motor memories.

## Introduction

The dopamine system and the basal ganglia play unique roles in motor control and motor learning.^1,2^ The D1 and D2 dopamine receptors are expressed in two distinct pathways: the D1 (direct) and D2 (indirect) pathways.^3,4^ Despite their well-recognized roles in facilitating and inhibiting movement,^5–7^ how each of these pathways contributes to learning processes and how they are involved in memory storage, particularly under pathological conditions, remain largely underexplored.

In Parkinson’s disease (PD), degeneration of dopamine neurons in the substantia nigra leads to significant disruptions in the balance between the D1 (direct) and D2 (indirect) pathways.^8,9^ We have previously reported that in addition to acute motor impairment caused by dopamine loss or dopamine receptor blockade, the impaired motor behavior is contributed largely by motor experience-dependent gradual deterioration of motor performance in the absence of dopamine receptor activation, a “use it and lose it” phenomenon we refer to as aberrant inhibitory motor learning.^10–12^

Aberrant Inhibitory learning reminds us of the concept of extinction learning; both processes are experience dependent and task specific, and lead to a decline in previously acquired responses. In extinction learning, the formation of extinction memory doesn’t necessarily erase the original memory; rather, it superimposes a new memory that opposes the behavioral level expression of the initial memory (For review, see ^13,14^). The similarities between extinction learning and aberrant inhibitory learning prompted us to ask: does aberrant inhibitory learning erase normal motor memory? In other words, are normal memory and aberrant inhibitory memory stored independently or even in separate anatomical pathways?

Our earlier studies suggest that aberrant inhibitory learning is mediated by the D2 (indirect) pathway.^10^ We therefore examined if normal memory and aberrant inhibitory memory are stored independently in the D1 (direct) and D2 (indirect) pathways respectively. We used behavioral designs to specifically probe normal motor learning versus aberrant inhibitory motor learning, and recorded GCaMP signals from D1 and D2 receptor-expressing dSPNs and iSPNs. In order to use manipulations to selectively impair, therefore dissociate, normal learning and aberrant inhibitory learning, we used a genetic approach by deleting the *Ythdf1* gene in dSPNs or iSPNs. Many studies have shown that new protein synthesis is required for long-term memory formation. YTHDF1 is an m^6^A RNA methylation reader protein, a specific RNA-binding protein that recognizes and binds to m^6^A modified mRNAs and facilitates their translation.^15^ It has been shown that it works with FMRP and plays a significant role in synaptic plasticity, learning and memory.^16,17^ Collectively, our behavioral, GCaMP recording and genetic data all support our hypothesis that normal memory and aberrant inhibitory memory are stored independently in the D1 (direct) and D2 (indirect) pathways respectively. Finally, we built a computational model based on activities of motor cortical neurons, D1 and D2 receptor-expressing neurons, and their interactions through the basal ganglia loops and validated the above hypothesis.

## Results

### Normal motor memory is preserved after aberrant inhibitory learning

We used our previously established approach to study the relationship between aberrant inhibitory learning and normal motor learning in wild type (WT) mice and Pitx3 mutant mice using an accelerating rotarod task (Figure 1A). Consistent with our previous report, WT mice trained on rotarod under dopamine antagonists treatment showed impaired behavior even after the washout of the drug (Figure 1B, two-way ANOVA on day 8-9, group effect, F (1, 13) = 13.25, p = 0.0030; group x time interaction F (1, 13) = 7.829, p = 0.0151). In the experiment, the mice (red group) during aberrant inhibitory learning showed stereotyped immobile behavior on the rotarod. This immobile behavior persisted when we re-exposed them to the rotarod task even after the dopamine antagonists were washed out (probe phase).

**Fig 1.**
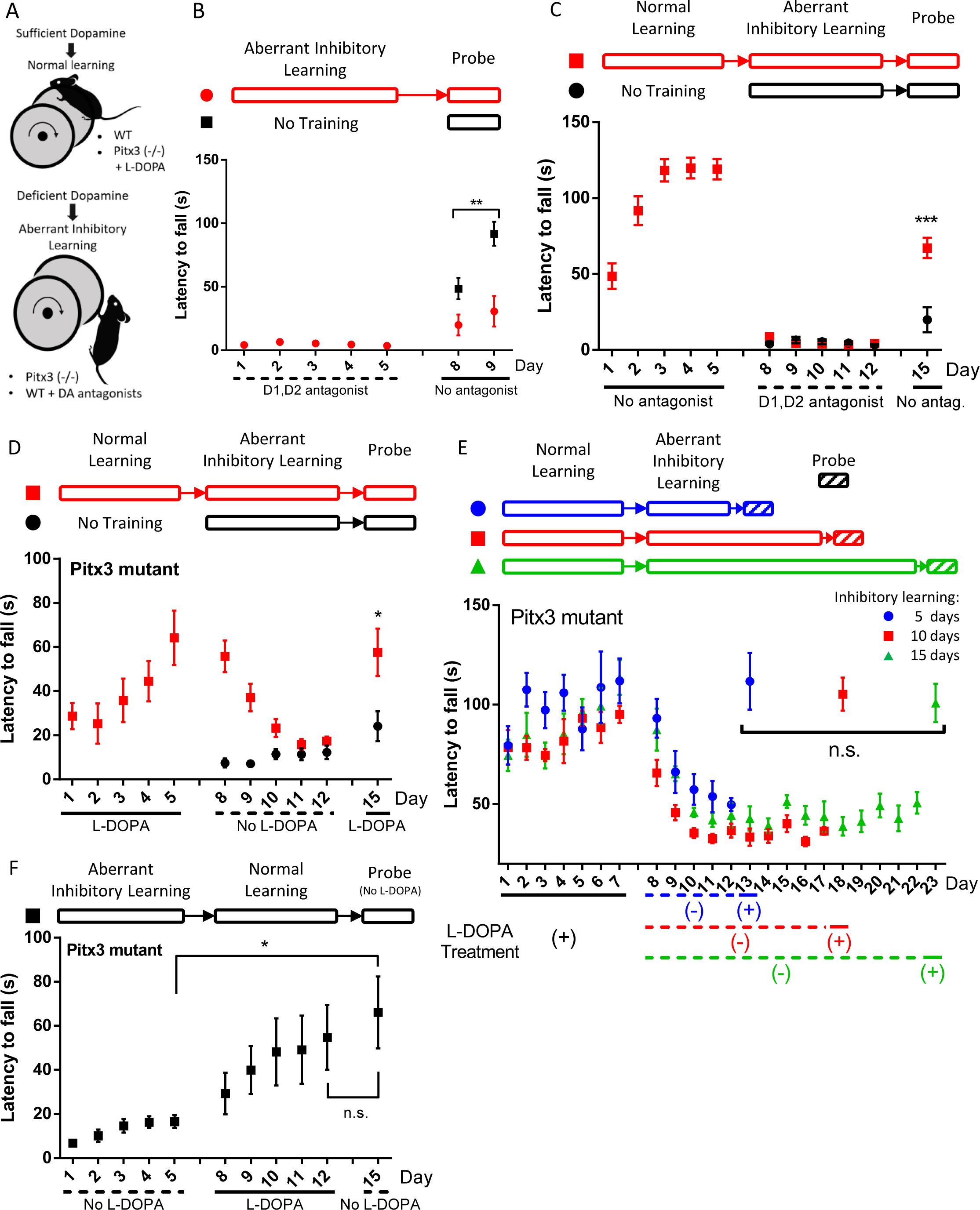
Normal memory was preserved after aberrant inhibitory learning, whereas inhibitory memory was reversed by normal learning. (A) Experimental designs for normal motor learning (WT mice without drug treatment or Pitx3 mutant mice treated with L-DOPA) and inhibitory motor learning (WT mice treated with dopamine antagonists or Pitx3 mutant mice without L-DOPA treatment). (B) Dopamine antagonist cocktail (SCH22390 and Eticlopride) treatment induced long-term impairment even after drug washout. Two-way ANOVA on day 8-9, group effect, F (1, 13) = 13.25, p = 0.0030; group x time interaction F (1, 13) = 7.829, p = 0.0151, n = 8 for each group. (C) In WT mice (red group) with previous normal learning experience showed significantly better performance in probe phase compared to the control (black) group. t test on day 15, p=0.0006, n = 7 for black group, n = 8 for red group. (antag., antagonist) (D) In Pitx3 mutant mice (red group) with previous normal learning experience showed significantly better performance in probe phase than the control (black) group, indicating preserved normal memory. t test on day 15, p=0.0251, n = 6 for each group. (E) In Pitx3 mutant mice, normal memory was equally preserved after 5, 10 or 15 days of inhibitory learning. One-way ANOVA of rotarod performance on L-DOPA treated days between 13 (blue), 18 (red) and 23 (green) day, F (2, 17) = 0.255, p = 0.778. n = 6 for the blue group with 5 days of inhibitory learning; n = 7 for the red group with 10 days of inhibitory learning; n = 7 for the green group with 15 days of inhibitory learning. (F) Pitx3 mutant mice showed sustained improvement from aberrant inhibitory learning after normal learning experience, indicating inhibitory memory was reversed. One-way ANOVA between day 5, 12, and 15, day effect F (2, 21) = 4.124, p = 0.0308. Post-hoc Tukey HSD test, Day 15 vs Day 5, p = 0.031, day 15 vs day12, p = 0.794, n= 8. All data represents mean ± SEM. *, p<0.05; **, p<0.01; ***, p<0.001. n.s., not significant.

To investigate whether inhibitory motor learning erases normal motor memory, we designed a three-phase motor learning paradigm involving motor skill learning on an accelerating rotarod using WT mice (Figure 1C, top). In the normal learning phase, only one group was trained on rotarod to acquire normal motor memory, while the other group received no training. In the aberrant inhibitory learning phase, both groups were treated with dopamine D1 (SCH22390) and D2 (Eticlopride) antagonist cocktails and trained on the rotarod. This procedure induced aberrant inhibitory learning in both groups and they both showed a performance deficit as expected. In the probe phase, both groups were tested under no drug condition on rotarod. This three-phase experiment allowed us to test if normal memory is preserved after inhibitory learning. To our surprise, we found that during the probe phase, the group with previous normal learning experience showed significantly better performance than the control group (Figure 1C and S1A-C; 1C, t test on day 15, P=0.0006). These data suggest that normal motor memory acquired before inhibitory learning is still preserved after inhibitory learning.

Next, we tested the same hypothesis using the Pitx3 deficient mutant mice that lack the nigrostriatal dopaminergic pathway throughout development.^18^ The experiment was similarly designed with three-phases (Figure 1D, top). One group of Pitx3 deficit mice went through normal learning (L-DOPA treatment), aberrant inhibitory learning (no L-DOPA) and probe phase (L-DOPA treatment). The second group of Pitx3 deficit mice went through only aberrant inhibitory learning and the probe phase. In normal learning phase, consistent with what we reported before,^11^ L-DOPA treated Pitx3 deficit mice successfully acquired the rotarod motor skill (Figure 1D). When both groups of mice were trained without L-DOPA during the aberrant inhibitory learning phase, the mice with normal learning experience showed a gradual decline of the motor performance, and the control mice showed low performance. Both groups reached the same level of low performance at the end of the aberrant inhibitory learning phase. In the probe phase, both groups were tested on rotarod again under L-DOPA condition. Remarkably, the group with previous normal learning experience showed significantly better performance than the control group (Figure 1D and S1D-F; 1D, t test on day 15, P=0.0251). These data indicate that the normal memory is still preserved in the Pitx3 deficit mice even after aberrant inhibitory learning.

To rigorously assess the durability of the preserved normal memory in Pitx3 deficit mice, we used the three-phase behavior design with different lengths of aberrant inhibitory learning (Figure 1E, top). All three groups of Pitx3 mutant mice were trained with L-DOPA in the normal learning phase to form the normal motor memory, and then they went through 5, 10, or 15 days of rotarod training without L-DOPA (aberrant inhibitory learning phase). Next, they were tested under L-DOPA condition to determine the extent to which normal memory had been retained (probe phase). Surprisingly, we found that all three groups showed similar levels of recovery in probe phase (Figure 1E and S1G-H; 1E, one-way ANOVA on probe phase, F (2, 17) = 0.255, P = 0.778). This observation suggests that the preservation of normal motor memory in these models is resilient to extended periods of dopaminergic deficiency and aberrant inhibitory learning.

Since normal memory is preserved after the aberrant inhibitory learning process, is the inhibitory memory also resistant to normal learning process? To test that, we designed another three-phase experiment using Pitx3 deficit mice. Mice were trained on an ‘aberrant inhibitory learning - normal learning - probe’ schedule (Figure 1F, top). The first two phases were intended to induce aberrant inhibitory memory and normal memory respectively. As expected, mice performance improved over days during the normal learning phase. Then mice were tested without L-DOPA on the probe phase. If the aberrant inhibitory memory was not reversed by the normal learning experience, we expected to see a similar performance between the aberrant inhibitory learning phase and the probe phase. To our surprise, the rotarod performance during probe phase did not show a sharp drop to the performance level of aberrant inhibitory learning (Figure 1F and S1I-J; One-way ANOVA between day 5,12, and 15, day effect p= F (2, 21) = 4.124, P = 0.0308; post-hoc Tukey HSD test, Day 15 vs Day 5, p = 0.031). On the contrary, the rotarod performance without L-DOPA treatment (probe phase) was maintained at a similar level as the end of normal learning phase (Figure 1F and S1I-J; post-hoc Tukey HSD test, day 15 vs day12, p = 0.794). These data suggest that inhibitory memory can be reversed by normal learning.

### dSPN and iSPN striatal neuron activities during normal and aberrant inhibitory learning

To examine if the preserved normal memory after aberrant inhibitory learning is reflected in activities of direct (dSPN) or indirect (iSPN) spiny projection neuron in the dorsal striatum, and to further characterize the neuronal activities during normal and aberrant inhibitory learning, we performed *in vivo* fiber photometry recording of Ca2+ activities during our rotarod motor learning paradigm. Previous studies had shown that dorsal striatum is important for rotarod motor learning.^19–22^ We injected a Cre-dependent GCaMP6m AAV9 into the dorsal striatum in either D1-Cre mice or A2a-Cre mice (Figure 2A, B) to express the calcium activity sensor in either the dSPN or iSPN respectively. We used the three-phase design described above (Figure 2C).

**Fig 2.**
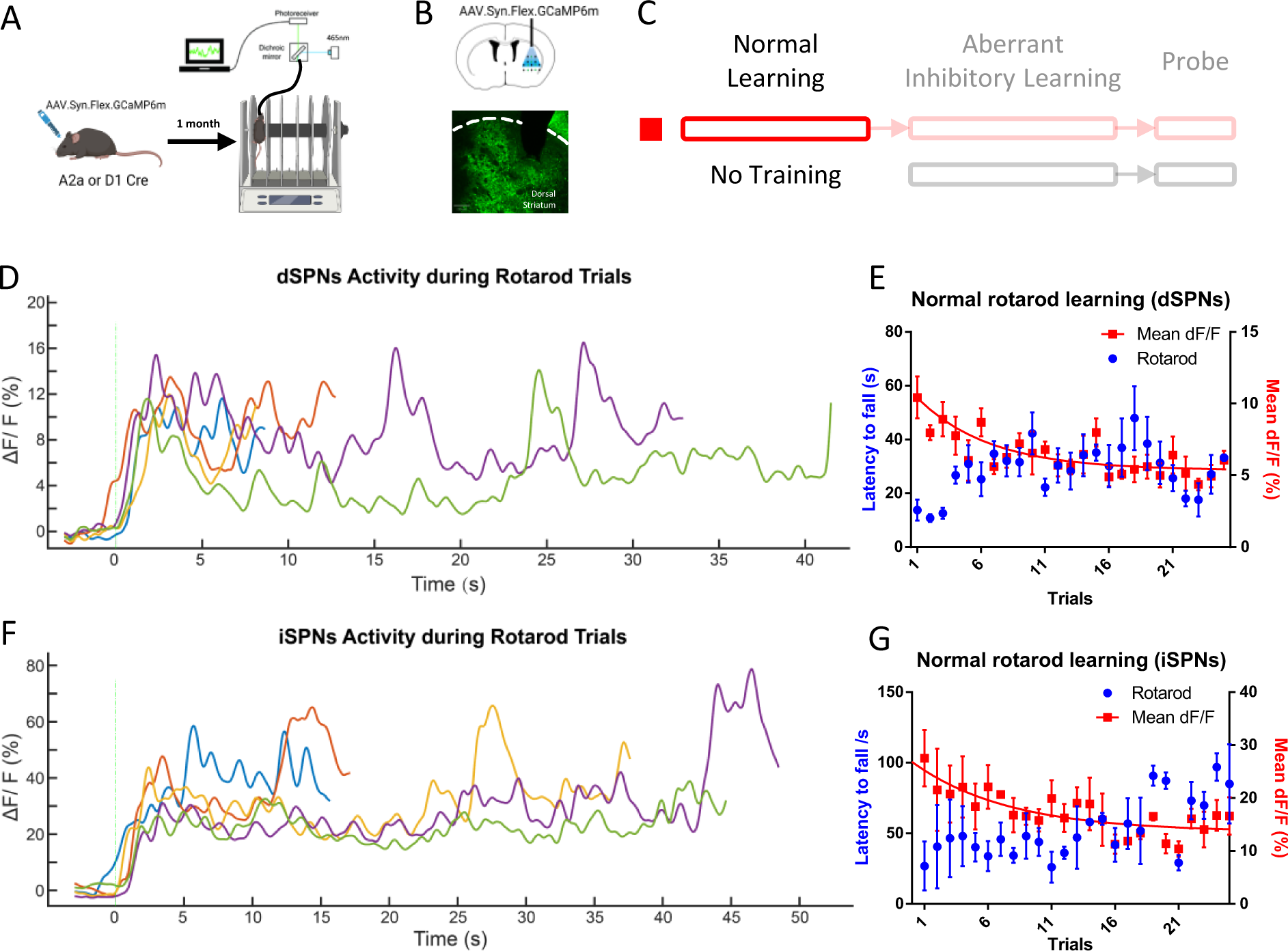
Fiber photometry recording of dSPNs and iSPNs in dorsal striatum during normal motor learning. (A) Experimental design for fiber photometry recording of Ca^2+^ activity in dorsal striatum during rotarod motor learning task. AAV-mediated dSPNs and iSPNs expression of GCaMP6m was achieved after local injection of Cre recombinase-dependent AAV into the dorsal striatum of D1-Cre and A2a-Cre transgenic mice respectively. (B) Top, schematic of injection site and optical fiber placement (black bar) for the GCaMP6m Ca^2+^ sensor. Bottom, fluorescence image showing GCaMP6m (green). White dashed line, border of striatum. Fiber placed between the tissue gap of dashed lines. Fiber placement position (AP +0.7, ML +2.25, depth 2.6mm). (C) The three-phase experimental design used for fiber photometry recordings. (D-G) Fiber photometry recordings during normal motor learning. (D) Representative traces of Ca^2+^ signal in dSPNs during rotarod test. Each trace is a single trial, with green dash line indicating the beginning of a trial. (E) Mean Ca^2+^ activity in dSPNs throughout the rotarod motor skill acquisition phase, n = 4. One-phase exponential decay in mean Ca2+ signal following rotarod training, fitted using the equation Y=(10.39 - 5.35)*exp(-0.169*X) + 5.35, half-life = 4.1, time constant = 5.92, R² = 0.28. Mean ± SEM. (F) Representative traces of Ca^2+^ signal in iSPNs during rotarod test. (G) Mean Ca^2+^ activity in D2 striatal neurons throughout the rotarod motor skill acquisition phase, n = 3. One-phase exponential decay in mean Ca2+ signal following rotarod training, fitted using the equation Y=(26.85 - 13.61)*exp(-0.13*X) + 13.61, half-life = 5.25, time constant =7.58, R² =0.24. Mean ± SEM.

During the normal motor learning phase, we found that both dSPN and iSPN calcium signals showed an increase at the beginning of the rotarod trial and stayed elevated throughout the trial (Figure 2D, F). Interestingly, as the mice learned the rotarod over several days, the average amplitude of the Ca2+ signal in both dSPN and iSPN gradually decreased (Figure 2E, G; 2E, One-phase exponential decay, Y=(10.39 - 5.35)*exp(-0.169*X) + 5.35, half-life = 4.1, time constant = 5.92, R² = 0.28; 2G, One-phase exponential decay, Y=(26.85 - 13.61)*exp(-0.13*X) + 13.61, half-life = 5.25, time constant =7.58, R² =0.24). This seems to suggest both the direct and indirect pathways are involved in normal motor learning. However, it is important to keep in mind that both direct and indirect pathways could change during motor learning, but they are not necessarily the mechanisms underlying motor learning.

We next studied the neuronal activities during aberrant inhibitory learning phase (Figure 3B, and 3C-J), where we treated mice with a dopamine antagonist cocktail (SCH22390 and Eticlopride). The mice under dopamine antagonists treatment had stereotyped immobile behavior on the rotarod, and this behavior persisted when the mice were re-exposed to the rotarod context even after the dopamine antagonists were washed out. To understand the neuronal activity underlying the immobile behavior (aberrant inhibitory learning), we divided the trials into short (<5s, mostly immobile behavior) and long (>10s) trials. When we examined the distribution of trial length in the two groups, we found that the group with previous normal learning experience had fewer short trials compared with the group not previously trained (Figure 3C, G). This is true for the later probe phase as well (see below and Figure 3K, O). In studying fiber photometry signals during the aberrant inhibitory learning phase, we aligned the start of the trial between long and short trials to understand the neuronal activity that contributed to the short trials. We found that there is no difference between short and long trials in the dSPN signal, in either group (Figure 3D, E). This observation is further confirmed when we compared the mean Ca2+ signal of the first 2 seconds of the trials. Neither trial length nor group factor is significant in a two-way ANOVA test (Figure 3F. Two-way ANOVA, trial length factor F (1, 92) = 0.088, P = 0.77; training experience factor F (1, 92) = 0.030, P=0.86). Next, we looked at fiber photometry signals in the indirect pathway during the aberrant inhibitory learning phase. There is no difference between short and long trials in the group with previous normal learning experience (Figure 3H). However, we found that in the short trials of the control group, iSPN signal sharply increased after the beginning of the trial (Figure 3I). When comparing the mean signal of the first 2s of the trial, the iSPN signal in the short trial of the control group is significantly higher than the long trials, and it’s also significantly higher than the short trials in the (red) group with previous normal training (Figure 3J. Post hoc Tukey HSD test: ‘black >10s’ vs ‘black <5s’, P=0.001; ‘red <5s’ vs ‘black <5s’, P=0.002). However, the group with normal learning experience earlier seemed more protected from such an effect. These data suggest that higher iSPN signal is associated with poor performance during aberrant inhibitory learning, which is consistent with the idea of a hyperactive indirect pathway during aberrant inhibitory learning, although it is conceivable that normal activities of iSPN and appropriate level of inhibition are necessary for normal learning as well.

**Fig 3.**
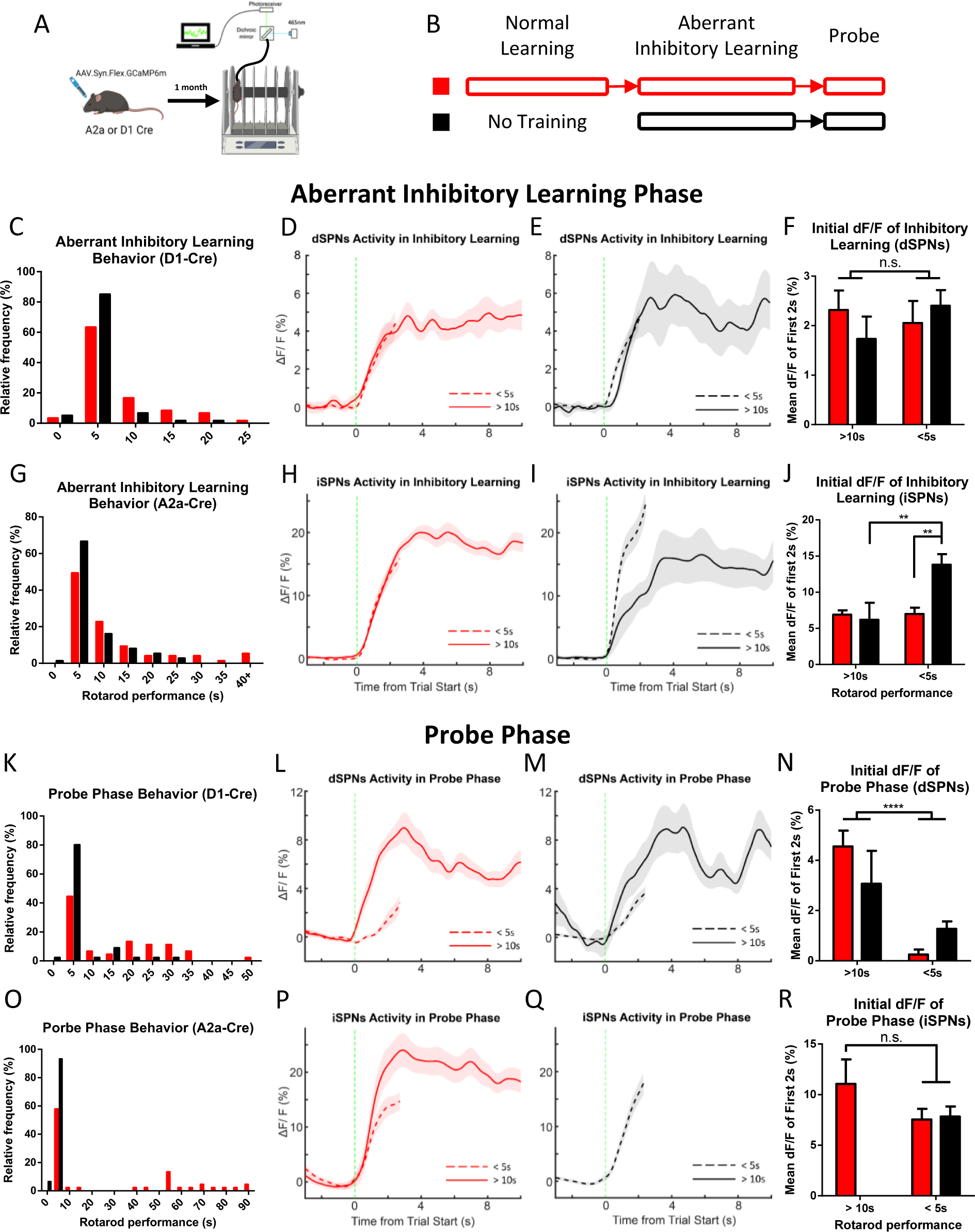
Preservation of normal memory after aberrant inhibitory learning was reflected in the activities of dSPNs. (A) Fiber photometry recording setup. (B) Experimental design for fiber photometry recording of Ca^2+^ activity in dorsal striatum during aberrant inhibitory learning and probe phase (Normal learning phase is shown in figure 2). Red group was trained in normal learning phase while black group did not receive normal motor learning. (C-J) Fiber photometry recordings during aberrant inhibitory learning phase. (C) Histogram of rotarod behavior during the aberrant inhibitory learning phase of mice used in dSPNs recordings. (D) Mean dSPNs Ca^2+^ signal of the red group (with previous normal learning experience) during aberrant inhibitory learning. Dashed line, < 5s trials; solid line, > 10s trials. Mean ± SEM. (E) Same as (D) but for the black group with no previous normal learning experience. (F) Comparing mean dSPNs Ca^2+^ signal of the first 2s of the aberrant inhibitory learning phase between red and black groups (with or without previous normal learning experience). There was a total of 43 trials from 3 red group mice and 53 trials from 3 black group mice. Two-way ANOVA, trial length factor F (1, 92) = 0.088, p = 0.77; group factor F (1, 92) = 0.030, p=0.86. Mean ± SEM. (G-J) Same as (C-F) but for iSPNs recordings. (I) In the black group without previous normal learning, higher iSPNs activities predict shorter trials. Mean ± SEM. (J) Comparing mean iSPNs Ca^2+^ signal of the first 2s between red and black groups. There was a total of 57 trials from 3 red group mice and 52 trials from 3 black group mice. Two-way ANOVA, trial length factor F (1, 105) = 7.93, p=0.0058; group factor F (1, 105) = 5.01, p = 0.027; Interaction F (1, 105) = 7.526, p=0.0072. Post hoc Tukey HSD test: ‘black group >10s’ vs ‘black group <5s’, Q statistic = 5.55, p=0.001; ‘red group <5s’ vs ‘black group <5s’, Q statistic = 5.24, p=0.002. (K-R) Same as (C-J) but for the probe phase of the fiber photometry experiment. (L) In the red group with previous normal learning experience, lower dSPNs activities predict shorter trials. Mean ± SEM. (M) In black group with no previous normal learning experience, lower dSPNs activities also predict shorter trials. Mean ± SEM. (N) Comparing mean dSPNs Ca^2+^ signal of the first 2s between red and black groups (with or without previous normal learning experience). There was a total of 43 trials from 3 red group mice and 43 trials from 3 black group mice. Two-way ANOVA, trial length factor F (1, 82) =28.51, p<0.0001; group factor F (1, 82) =0.168, p=0.68. Mean ± SEM. (R) Comparing mean iSPNs Ca^2+^ signal of the first 2s between red and black group. There was a total of 42 trials from 3 red group mice and 45 trials from 3 black group mice. Unpaired t test between >10s and <5s, p=0.08. Mean ± SEM. *, p<0.05; **, p<0.01; ***, p<0.001; ****, p<0.0001. n.s., not significant.

After dopamine antagonist induced aberrant inhibitory learning, all the groups are tested under the drug-free condition (Figure 3B and 3K-R, probe phase). Again, when we examined the distribution of trial length in different groups, we found that trials in the group with previous normal learning showed less distribution in the short trials compared with the control group, consistent with our behavior studies (Figure 3K, 3O and S1C). Similar to the above analysis in the aberrant inhibitory learning phase, we divided the trials into long (>10s) and short (<5s) trials. In the fiber photometry signal, we found that the dSPN signal in short trials showed a slower ramp compared with the long trials (Figure 3L, M). This observation was consistent in both groups with or without previous normal learning experience, and was further confirmed by the significantly higher average signal of the first 2 seconds of the long trials (Figure 3N. Two-way ANOVA, trial length factor F (1, 82) =28.51, P <0.0001). The fiber photometry signals in the indirect pathway showed a similar trend of difference between long and short trials but were not statistically significant (Figure 3R. Unpaired t test between >10s and <5s trials, P=0.08). These data suggest that a faster activity ramp in the D1 pathway is associated with better performance. Given that the group with previous normal learning has more long trials with a faster activity ramp in the D1 pathway compared with the control group, we conclude that previous normal training experience before aberrant inhibitory learning induced higher dSPN activity during the drug-free probe phase, potentially an underlying mechanism for their better performance in the probe phase.

To examine whether there are similarities in fiber photometry signals between the normal learning phase and the probe phase, we performed principal component analysis (PCA) (Figure S2) using several features extracted from each individual trial in the two phases (when performance was not under drug influence). The photometry signal features we used in PCA include: signal peak rate, trial signal mean, trial standard deviation, signal mean during trial beginning, and signal mean during trial end. In analysis of the dSPN fiber photometry data, principal components 1 and 2 successfully explained 82.0% and 12.5% of the total variance respectively. When we plotted all the trials together using principal component 1 and 2, we observed a clustering pattern in which there is a clear separation between the normal learning phase and the probe phase data from the group without previous normal learning (Figure S2B). However, the probe phase data from the group with previous normal learning (recall of normal learning) overlaps with normal learning phase and with probe phase data from the group without previous normal learning, potentially suggesting that the group with previous normal learning showed features of both the normal memory and aberrant inhibitory memory during the probe phase. Similarly, we performed such PCA analysis on iSPN fiber photometry data, principal component 1 and 2 explained 84.0% and 10.9% of the variance respectively. However, when we plotted all the trials using PC1 and PC2, the iSPN data from the normal learning phase and the probe phase were mixed and did not show a clear clustering pattern (Figure S2C).

### Dissociation of D1 pathway-dependent normal learning and D2 pathway-dependent inhibitory learning

Our earlier studies suggest that inhibitory motor learning is mediated by the D2 (indirect) pathway.^10^ The present fiber photometry data suggest that activities in the D1 (direct) pathway is more important for normal motor learning, which is in agreement with previously reported optogenetic inhibition studies.^20^ To further investigate the roles of D1 and D2 pathways in normal and aberrant inhibitory motor memory, we designed a double dissociation experiment utilizing conditional *Ythdf1* gene deletion. Previous studies showed that *Ythdf1* gene deletion impaired new protein synthesis, synaptic plasticity in the hippocampus and hippocampus-dependent learning.^17^ To confirm the role of YTHDF1 in regulating new protein synthesis in the striatum, we measured the new protein synthesis rate using click chemistry technology in primary striatal neuronal culture.^23–25^ Specifically, we incubated the striatal neuronal culture with a methionine analog HPG in a methionine free medium to label the newly synthesized protein, and later tagged the HPG with fluorophore using click chemistry reaction for visualization. We found that the fluorescence intensity was much higher in the baseline group compared with a negative control group, where we treated cells with cycloheximide (CHX), a protein translation inhibitor (Figure 4A, B and S3A. One-way ANOVA, F(2,99)=97.7, P<0.0001; post hoc Tukey HSD test, ‘CHX’ vs ‘Baseline’, Q statistic = 5.54, P=0.001). After validation of the method, we treated cells with forskolin to activate adenylyl cyclase and its downstream signaling as an approach to activate intracellular protein synthesis. We saw that forskolin treatment significantly increased HPG signal, which indicated higher protein synthesis rate (Figure 4A, B and S3A. Post hoc Tukey HSD test ‘Baseline’ vs ‘Forskolin’, Q statistic = 12.88, P=0.001). Surprisingly, when we performed the same procedure in the striatal neurons derived from *Ythdf1* knockout, they did not show such an increased protein translation when cells are treated with forskolin (Figure 4A, C and S3B. T-test ‘Baseline’ vs ‘Forskolin’, p<0.0001). On the contrary, striatal neurons from the *Ythdf1* knockout showed a much higher baseline protein translation rate than the neurons in WT. This result suggests that *Ythdf1* KO striatal neurons are impaired in activating protein translation in response to extracellular stimuli. While it is conceivable that cells that do not respond to elevated activities by elevating protein synthesis rate need to have a higher baseline protein translation rate in order to sustain normal cellular functions with sufficient protein translation activities, the consequences of elevated baseline protein translation rate are worth investigating in the future. Of note, the higher protein synthesis rate caused by stimulation is far lower than the maximum limit of our assay, ruling out a ceiling effect.

**Fig 4.**
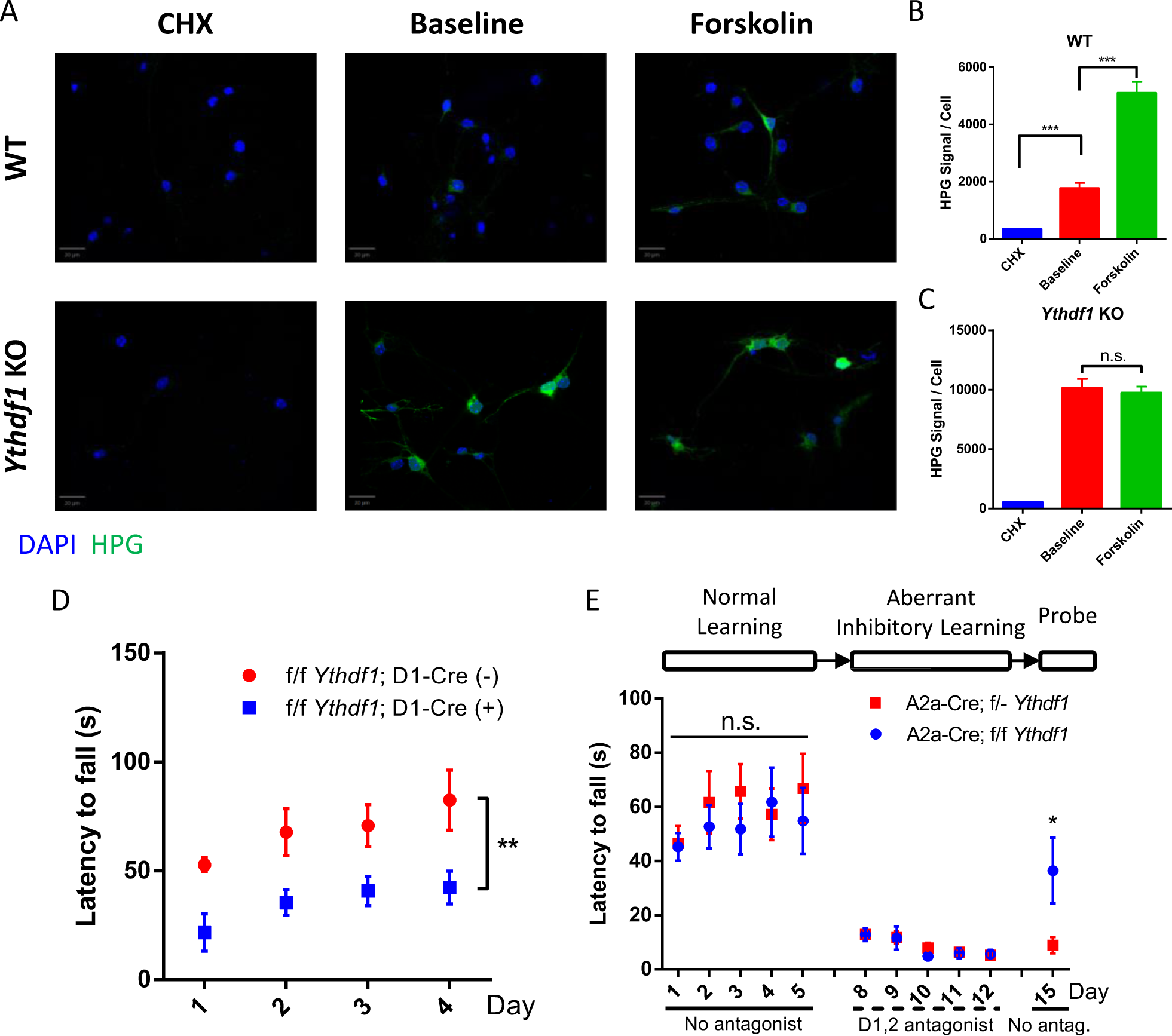
*Ythdf1* Gene deletion experiments showed double dissociation and suggested that normal learning and inhibitory learning are mediated mainly by the D1 and D2 pathways respectively. (A) Representative merged images of click chemistry experiment measuring protein synthesis rate in WT and *Ythdf1* KO mice. Blue, DAPI staining; green, HPG tagged newly synthesized protein. Scale bar, 20 μm. CHX, Cycloheximide. (B) Quantification of newly synthesized protein during CHX, baseline and forskolin treatment in WT striatal neurons. One-way ANOVA, F(2,99)=97.7, P<0.0001; post hoc Tukey HSD test, ‘CHX’ vs ‘Baseline’, Q statistic = 5.54, p=0.001; ‘Baseline’ vs ‘Forskolin’, Q statistic = 12.88, p=0.001. (C) Quantification of newly synthesized protein during CHX, baseline and forskolin treatment in *Ythdf1* KO striatal neurons. Student t-test ‘Baseline’ vs ‘Forskolin’, p=0.6927. n= 36 for CHX, and Forskolin group, n= 30 for HPG group. Each group contains 3 replicates. (D) Conditional gene deletion of *Ythdf1* in D1-Cre mice led to an impairment in the normal motor learning paradigm. Two-way ANOVA, group effect F(1, 12) =11.23, p= 0.0058. Group x time interaction F(3, 36) = 0.2895, p=0.8327. n= 7 for each group. (E) Conditional gene deletion of *Ythdf1* in A2a-Cre mice led to similar normal motor learning but more protection against aberrant inhibitory motor learning compared to the control group. Two-way ANOVA on day 1-5, group effect F(1, 9) =0.2586, p=0.6233. Group x time interaction F(4, 36) = 0.8894, p=0.4802. t-test on day 15, p=0.0398. n(A2a Cre x f/f *Ythdf1*) = 5. n(A2a Cre x f/- *Ythdf1*) = 6. All data represent mean ± SEM. *, p<0.05; **, p<0.01; ***, p<0.001. n.s., not significant.

After validating the role of YTHDF1 in regulating protein synthesis in the striatum. We deleted *Ythdf1* in the D1 or D2 pathway. We hypothesize that impaired new protein synthesis in dSPN or iSPN should impair D1 or D2 pathway dependent learning respectively. We first tested whether the D1 pathway is important for normal motor learning using D1 neuron specific *Ythdf1* gene deletion (D1-Cre x floxed-*Ythdf1)* and control mice of 6 months. We found that rotarod performance during normal learning was significantly impaired in mutant mice compared to controls (Figure 4D. Two-way ANOVA, group effect F(1, 12) =11.23, P= 0.0058; group x time interaction F(3, 36) = 0.2895, P=0.8327). Because the normal memory formation was already impaired in D1-Cre x floxed-*Ythdf1* mice, this prevented us from further testing if they were impaired in inhibitory learning.

Next, we investigated the involvement of the D2 pathway of the basal ganglia in normal and inhibitory learning. Using A2a-Cre mouse line crossed with floxed-*Ythdf1*, we generated mice with conditional deletion of *Ythdf1* in D2 dopamine receptor-expressing cells. We found that 6-month-old mutant mice showed no difference during normal learning compared with the controls (Figure 4E. Two-way ANOVA on day 1-5, group effect F(1, 9) =0.2586, P=0.6233; group x time interaction F(4, 36) = 0.8894, P=0.4802), suggesting that new protein synthesis in the D2 pathway is not directly involved in normal motor learning. To investigate whether the aberrant inhibitory learning is affected, we trained mice on rotarod under dopamine D1 & D2 antagonist cocktails, and then tested both groups without drug injection (Figure 4E, top). We found that conditional knockout of *Ythdf1* in the D2 pathway showed significantly better performance (less inhibitory learning) during the probe phase (Figure 4E, S3C. 4E, unpaired t-test on day 15, P=0.0398), suggesting that inhibitory learning is mediated through the basal ganglia D2 pathway. Because YTHDF1 regulates protein synthesis rate, this result suggested that manipulating protein synthesis rate in the D2 pathway may prevent the formation of aberrant inhibitory learning under dopamine deficient conditions.

### Computational Model supports preserved D1 normal memory after inhibitory learning

To recapitulate and understand how dopamine regulates the acquisition, impairment, and recovery of motor skill, we established a computational model based on the classical “cortico-basal ganglia-cortical loop” architecture (see Supplemental Information for the model description). In the model, the activity of motor cortex that controls the rotarod task is modulated by a positive feedback loop via striatal D1 neurons (i.e. the direct pathway), as well as a negative feedback loop via D2 neurons (i.e. the indirect pathway) (Figure 5A). The core hypothesis underlying the model is the “cAMP-protein synthesis-memory consolidation” process (Figure 5B). We hypothesized that by regulating protein synthesis through dopamine receptor coupled cAMP pathway, consolidation of memory/plasticity is affected accordingly.^26,27^ In the model, increased levels of dopamine activate both D1 and D2 dopamine receptors, which induces LTP and LTD in the equivalent connection weights of the positive and negative feedback pathways, respectively. Consequently, this facilitates the formation of normal memory but diminishes inhibitory memory.^28^ Further, the level of dopamine regulation is proposed to be proportional to the behavioral prediction error of the animal, which is anticorrelated to the task performance.^29^

**Fig 5.**
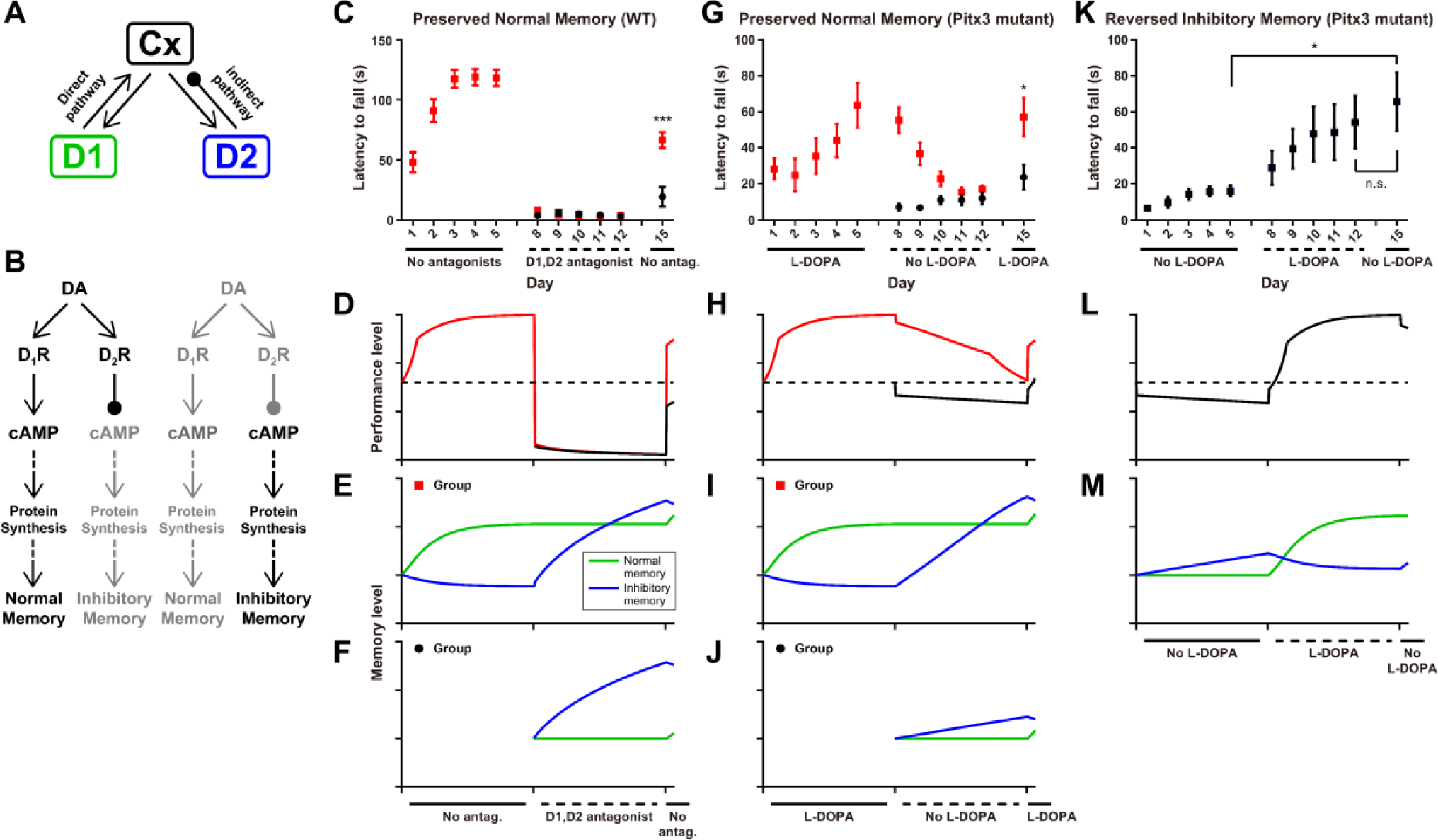
Modeling dopamine effects on mice rotarod performance under various experimental conditions. (A) Schematic of the “cortico-basal ganglia-cortical loop” model containing direct and indirect feedback pathways. “Cx” reprsents the motor cortex. (B) Schematic of the “cAMP-protein synthesis-memory consolidation” model of dopamine regulation on long-term plasticity of memories. (C) Observed rotarod performance data in “normal learning – aberrant inhibitory learning – probe” (red) and “aberrant inhibitory learning – probe” (black) experimental design. (D) Predicted rotarod performance by the model. (E - F) The weights of normal memory and aberrant inhibitory memory in the model. (G − J) Similar to (C − F) but in Pitx3 mutant mice. (K − M) Similar to (C − E) but in Pitx3 mutant mice using the “aberrant inhibitory learning - normal learning - probe” design.

We next tested if our computational model can recapitulate our behavior data. We found that the behavioral performance of WT mice in the experiments (Figure 5C) and model (Figure 5D) were well matched. Notably, during the aberrant inhibitory learning phase of the experiment, the application of dopamine antagonists blocks dopamine receptors, leading to an immediate drop of motor cortex activity and performance (Figure 5D). As training continues, the receptor blockade halts normal memory learning, but promotes the aberrant inhibitory memory learning (Figure 5E). During the probe phase where antagonists are removed, the normal memory is well preserved for the group with previous normal learning, but much weaker in the group without previous normal learning (Figure 5E vs 5F). This difference in the level of normal memory explains the dramatic performance difference in the two groups right after drug removal when the probe phase begins (Figure 5D). The recovery of normal memory may not be 100% in WT or in Pitx3 mutant mice treated with L-DOPA (see below); it is affected by synaptic strength changes in both the direct and indirect pathways. We expect that many factors (e.g., dopamine receptor sensitivity, age, genetic background etc.) could contribute to the variation in the recovery strength.

Our model also recapitulates our behavior findings using Pitx3 mutant mice (Figure 5G, 5H). In the Pitx3 mutant mice, normal memory is preserved while aberrant inhibitory learning is boosted during the “No L-DOPA” phase where the animals lack both endogenous and external dopamine (Figure 5I, 5J). One difference between the Pitx3 mutant mice and WT mice treated with antagonist is the smooth decay of motor cortical activity and performance over time during the aberrant inhibitory learning phase in Pitx3 mutant mice, rather than an instantaneous fall in WT mice treated with antagonist (Figure 5D vs 5H). One possible underlying mechanism is the adaptation of D1, D2 receptor activity to low dopamine level in the Pitx3 mutant mice (details in supplemental information).

Additionally, the model also captures the reversal of aberrant inhibitory memory in the ‘aberrant inhibitory learning-normal learning-probe’ schedule (Figure 5L, 5M). In the model, L-DOPA drives LTP and LTD of D1 and D2 pathways, respectively, leading to a boost of the normal memory and decay of the aberrant inhibitory memory (Figure 5M, “L-DOPA” phase). The changes in the relative strength of normal memory and aberrant inhibitory memory lead to improved performance level (Figure 5L, 5M), which would maintain even after the removal of L-DOPA before any updates by additional learning (Figure 5L, the second “No drug” phase).

## Discussion

Motor impairments in PD are often attributed to the lack of dopamine. However, our previously published studies^10–12^ as well as the present data indicate that the impaired motor behavior is contributed largely by motor experience-dependent gradual deterioration of motor performance in the absence of dopamine or dopamine receptor activation, a mechanism that we refer to as aberrant inhibitory motor learning. Although the opposing consequences of normal motor learning and aberrant inhibitory learning are manifested in apparent motor performance, at the circuit level, normal motor memory and aberrant inhibitory memory are stored in separate circuits, the former in the D1 (direct) pathway and the latter in the D2 (indirect) pathway. This is supported by our behavior results, fiber photometry recordings of GCaMP signals from dSPN and iSPN in the dorsal striatum, and genetic double dissociation of D1 pathway-dependent normal learning and D2 pathway-dependent aberrant inhibitory learning. It is also supported by a computational model based on activities of motor cortical neurons, D1 and D2 receptor-expressing neurons, and their interactions through the basal ganglia loops.

PD causes deteriorated motor control.^30,31^ In animal models, our behavior results here showed that even though the impaired motor behavior was encoded into long term memory, the normal memory learned under normal dopamine condition prior to the aberrant inhibitory learning is still preserved in the brain. These findings suggest that reactivating preserved normal memory could be therapeutic in PD. Indeed, in dopamine replacement therapy for PD, there are phenomena parallel to our findings in animal models. Long duration response is often observed in addition to the acute short duration response of the therapeutic effect.^32,33^ The long duration response is a gradual buildup of the therapeutic effect through many days until it reaches the maximum strength, which is similar to the reversal of aberrant inhibitory learning by normal learning in our data (Figure 1F). On the other hand, the gradual decay of long duration response, which typically lasts many days after cessation of dopamine replacement therapy, is similar to the aberrant inhibitory learning process in our data (second phase in Figure 1D), i.e., the experience-dependent gradual deterioration of motor performance in the absence of dopamine signaling. Surprisingly, even after complete decay of long duration response, the first dose of L-DOPA or dopamine agonist in PD patients can often cause a complete rebound to the maximum therapeutic effect, without going through the gradual reversal of aberrant inhibitory learning by normal learning, similar to immediate reactivation of normal motor memory in our data (Figure 1D).

Our fiber photometry studies give additional insights into the distinct roles played by D1 and D2 pathways in motor memory. A key observation is that high D1 pathway activities predict long rotarod trials whereas low D1 pathway activities predict short rotarod trials after aberrant inhibitory learning. This suggests that normal motor memory stored in the D1 (direct) pathway remains largely intact and can be potentially reactivated, despite the progression of motor impairments in PD. Of note, due to the technical limitation of fiber photometry, we could not attain calcium dynamics at cellular resolution. Future studies with more advanced imaging techniques will help understand the dynamics of normal and inhibitory memory engram during learning. Imaging technique with cellular resolution will also help us understand whether the changes we observe is due to somatic or non-somatic calcium activities, since a recent study showed that fiber photometry in striatum mostly reflect non-somatic changes.^34^

Both dopamine D1 and D2 receptors are coupled to Adenylyl cyclase 5 (AC5) and therefore the cAMP pathway. However, they have opposite effects on AC5 and protein kinase A (PKA) activity upstream of YTHDF1 and protein synthesis, with the D1 receptor activating while the D2 receptor inhibiting the cAMP pathway. It has been demonstrated that PKA activity in spiny projection neurons is dynamically influenced by dopamine differently in D1 versus D2 striatal neurons.^35,36^ With increased dopamine, PKA is activated in D1 neurons. In contrast, PKA is activated in D2 neurons in dopamine-deficient states. Elevated cAMP and activated PKA can affect downstream signaling including the expression of immediate early genes, CREB, and new protein synthesis, all implicated in memory consolidation.^27,37,38^ These mechanisms emphasize the essential role of elevated but not reduced cAMP that leads to new protein synthesis in memory consolidation, and may explain why the D1 pathway mediates normal motor learning and memory under normal dopamine whereas the D2 pathway mediates aberrant inhibitory motor learning and memory under dopamine deficiency. An alternative hypothesis to explain the distinct roles of the D1 versus D2 pathway in normal versus aberrant inhibitory learning/memory respectively is the receptor affinity hypothesis.^39^ D1 receptors have low affinity for dopamine, therefore they are not activated at the baseline condition, and they are more sensitive to increased dopamine release. In contrast, D2 receptors have high affinity for dopamine, therefore they are already activated at the baseline condition, and they are only sensitive to decreased dopamine release. This alternative hypothesis does not rely on the cAMP-new protein synthesis hypothesis. However, recent data challenges the D1 low affinity and D2 high affinity hypothesis.^40–44^ Our computational model effectively corroborates our empirical results, lending further support to the cAMP-memory consolidation hypothesis. This computational approach also provides a powerful tool for predicting and understanding the complex dynamics of memory consolidation in the basal ganglia under varying conditions, and potentially guiding the development of targeted therapeutic strategies.

Our study highlights the integral role of protein synthesis in memory consolidation within the basal ganglia in addition to its well characterized role in the hippocampus.^45–47^ The contrasting effects of *Ythdf1* knockout in the D1 versus D2 pathways suggest potential therapeutic targets for PD and other disorders. Most importantly, our result shows that conditional knocking out of *Ythdf1* in the D2 pathway did not affect normal learning process, but only mitigated the aberrant inhibitory learning, providing substantial therapeutic potential. By altering/modulating new protein synthesis pathways, it may be possible to enhance normal motor memory learning and mitigate the consolidation of aberrant inhibitory memories. This approach could lead to novel treatments that focus not only on symptomatic relief but also on the underlying synaptic neurobiological mechanisms of the disease which can potentially achieve long-lasting therapeutic effects.

Our findings also suggest compelling similarities between the aberrant inhibitory motor learning and the phenomenon of extinction learning. For example, in fear conditioning, current theories propose that extinction involves the formation of a new associative memory that inhibits the expression of the pre-existing memory, rather than erasing it.^14,48^ This newly formed ‘extinction memory’ affects the manifestation of the antecedent memory trace. Similarly, we have demonstrated that aberrant inhibitory motor learning, induced by dopaminergic deficits and motor experience, does not erase pre-existing normal motor memories. Despite the similarities, there are also distinct features in our data: 1) there is clear anatomical segregation of pathways underlying normal motor memory (D1 pathway) and aberrant inhibitory motor memory (D2 pathway). In extinction learning, no clear anatomical segregation has been reported; 2) aberrant inhibitory learning can be latent. When wild-type mice were treated with dopamine antagonists and trained on the rotarod, they apparently were not “learning” anything. However, their aberrant inhibitory learning was only revealed when they were trained again under no drug condition (Figure 1B). Even though latent extinction has been reported in the literature, it usually involves prevention of responses, and the mechanism is believed to be different from extinction learning.^49^ In addition to the similarities to extinction learning, the concept of aberrant inhibitory learning can potentially be applied to other D2 pathway dependent learning under normal physiology conditions, such as reversal learning and discrimination.^50,51^

In conclusion, we have demonstrated that normal motor memory is mostly stored in the D1 (direct) pathway whereas aberrant inhibitory motor memory is mostly stored in the D2 (indirect) pathway. The reactivation of either or both memories determines the apparent motor performance. These findings have important implications for novel therapeutic approaches in treating Parkinson’s disease by reactivating preserved normal memory, and in treating hyperkinetic movement disorders such as chorea or tics by erasing aberrant motor memories. The cAMP pathway and RNA binding proteins that facilitate new protein synthesis are important molecular targets to consider.

## Method

### Transgenic mouse Pitx3

#### mutant

Pitx3 (ak) mutant mice (Jackson Strain #:000942) exhibit an almost total loss of tyrosine hydroxylase-positive cells in the substantia nigra pars compacta, with a 90% decrease in dorsal striatal dopamine neuron at P0. The Pitx3 mutant mice are blind, yet this condition does not markedly influence their performance in the rotarod task used in the study.

#### Floxed *Ythdf1*, D1-Cre, A2a-Cre

Mice carrying a conditional removable *Ythdf1* allele (Ythdf1^f/f^) were crossed to a D1-Cre transgenic line (RRID: MMRRC-030989-UCD) or A2a-Cre transgenics line (RRID: MMRRC_036158-UCD) to selectively delete *Ythdf1* in D1 or D2 dopamine receptor expressing cells. All experiments were performed in both double transgenic mice (D1-Cre;Ythdf1^f/f^, A2A-Cre;Ythdf1^f/f^), and the respective control littermates.

## Mouse Behavior

### Rotarod

Mice in the task are 8-12 weeks old unless otherwise stated. A computer-controlled rotarod apparatus (Rotamex-5, Columbus Instruments, Columbus, OH) with a rat rod (7cm diameter) was set to accelerate from 4 to 40 revolutions per minute over 300 seconds, and recorded time to fall. Mice received 5 consecutive trials per session, 1 session per day. Rest between trials was approximately 30 seconds.

### Drug Administration

All drug injections were intraperitoneal at 0.01ml/gram of body weight. L-DOPA (3,4-dihydroxy-L-phenylalanine 25 mg/kg with 12.5mg/kg benserazide) was administered 1 hour prior to the start of each session. SCH 23390 at 0.1mg/kg and eticlopride at 0.16mg/kg were administered 30 minutes prior to experiments.

### Stereotaxic Surgery

All surgical procedures were performed using mice aged 12-16 weeks under sterile conditions. Mice were anesthetized using 2% isoflurane and placed in a stereotaxic frame. Skull was exposed and bregma - lambda was identified, hole was drilled above dorsal striatum (AP +0.7, ML +2.25), a guide needle was lowered 2.7mm DV, 400nL of AAV virus (Addgene Catalog # 100838, AAV9.Syn.Flex.GCaMP6m.WPRE.SV40) was delivered at a speed of 100nL/min, and allows for 7min to diffuse post injection before needle retraction. An optic cannula (MFC_400/430-0.66_5mm_MF1.25_FLT, Doric) was inserted into the injection site, 100μm above the viral delivery site. The cannula was then secured using surgical glue and dental cement.

### Fiber photometry

TDT-Doric system was used for fiber photometry studies, TDT RZ5P for signal driving and demodulation. This system was adept at delivering light at wavelengths of 405 nm and 465 nm, while monitoring at 525 nm through a specialized Doric minicube (FMC5_IE(400- 410)_E(460-490)_F(500-540)_O(580-680)_S, Doric). The received light was processed by a femtowatt photodetector (Newport Model 2151), which then channeled the signals to the RZ5P. We used distinct modulation frequencies to monitor signals based on calcium dependence. The 465 nm excitation light was calcium-responsive and modulated at 331Hz, while the 405 nm, an isosbestic calcium-independent control, was modulated at 211 Hz using LEDs and LED driver (Doric). Mice were tethered to a patch cord (0.48NA, 400 μm core diameter, Doric) with freely rotary joint and gimbal holder (Doric) for maximum freedom during movement. The TDT Synapse software was employed to interact with the RZ5P system, facilitating data logging, event timestamping via TTL loggers, and LED control. All data were analyzed in MATLAB with custom script, detailed code could be made available upon reasonable request. Briefly, first 5s recording was removed for opto-electro artifacts that might significantly affect the fitting parameters in the subsequent step. A smoothed 405nm signal was fitted to the 465nm signal using linear regression to obtain fitting coefficients during a-3s baseline period before every rotarod trial. Using the coefficients, we calculated the fitted 405nm and calculated normalized ΔF/F for the calcium fluorescence.

### Neuronal culture

Primary striatum neurons were cultured in 8-chambered coverglass systems (Cellvis C8-1.5H-N). Dissection was performed under a stereoscope utilized ice-cold 1x PBS, involving pia membrane removal and dorsal cortex dissection to expose the striatum. The dissected striatum tissues underwent enzymatic digestion with prewarmed Papain solution. After gentle chopping and incubation, the digested tissue was centrifuged, and cells were plated at a density of 0.04 million cells per well. Plating media transitioned to Neuromaintaining media after two hours. Medium maintenance involved replacing half the medium on day four and adding AraC to suppress gliogenesis. Subsequently, half the medium was regularly replaced every three days. Plating media included DMEM medium with 1% L-Glutamine, 1% penicillin–streptomycin, 0.8% Glucose, and 10% fetal bovine serum. Neuromaintaining media comprised Neurobasal medium with 1x B-27 supplement, 1x N2 supplement, 1% L-Glutamine, and 1% penicillin–streptomycin.

### Click chemistry

Methionine-free DMEM was prepared by adding 4mM glutamine 0.4mM cysteine (thermo scientific #J60573.14, #J63745.14) into customized DMEM (thermo fisher #21013024) and stored at 4C. HPG Alexa Fluor™ 488 kit was purchased from Thermo Fisher (#C10428). Cultured cells were gently washed with PBS and changed into methionine-free DMEM for 1-hour to decrease the intracellular methionine concentration. 5 μg/ml CHX and 10 μM Forskolin were added 10 minutes before adding HPG. Cells were added with a final concentration of 100 μM, HPG and incubated for 2 hours. Cells are washed with PBS and followed up with HPG labeling process described in protocol from thermo fisher. Cells are washed with PBS and incubated with MAP2 antibody (Sigma-Aldrich, Cat# M4403) for 2 hours at room temperature before the DNA staining step.

### Quantification and statistical analysis

Data are reported as mean ± SEM, and n represents the number of mice used per experiment unless otherwise stated. Statistical analyses were conducted in Graphpad. Statistical significance was assessed using a student’s t test or repeated-measures ANOVA for experiments that tracked behavior over time or repeated training, as well as fiber photometry experiments that compare different trials from different mouse groups. For significant findings after ANOVA, post-hoc Tukey’s HSD tests were used to identify specific group differences. The level of significance was set at p < 0.05.

### Computational model

Details are included in supplementary information.

### Author Contributions

K.W. and X.Z. conceived the project and experiments. P.Y., K.W., X.Z. and B.D. conceived the computational model. K.W. and Z.S. performed most of the experiments with help from L.M., K.W., V.Y., N.W. and J.B.. S.S. and D.M. helped with fiber photometry experimental design and data analysis. P.Y. built the computational model. K.W., P.Y. and X.Z. wrote the first draft of the manuscript while all authors contributed to the writing.

## Acknowledgements

We thank Benjamin Wang, Nabilah Sammudin, Nicholas LoRocco and Wenqin Fu for helpful discussions and comments on the manuscript. This work was supported by R01DA043361 (X.Z.), R01NS095374 (X.Z. and D.M.), Simons Foundation Collaboration on the Global Brain (B.D.), NIH 1U19NS107613-01 (B.D.), R01EB026953 (B.D.), and R01EY034723 (B.D.). Shared equipment grants from the University of Chicago Neuroscience Institute supported the shared fiber photometry, AAV injection, and histology facilities.

