## Supplemental info for "Opposing Motor Memories in the Direct and Indirect Pathways of the Basal Ganglia"

### Supplementary Figure 1

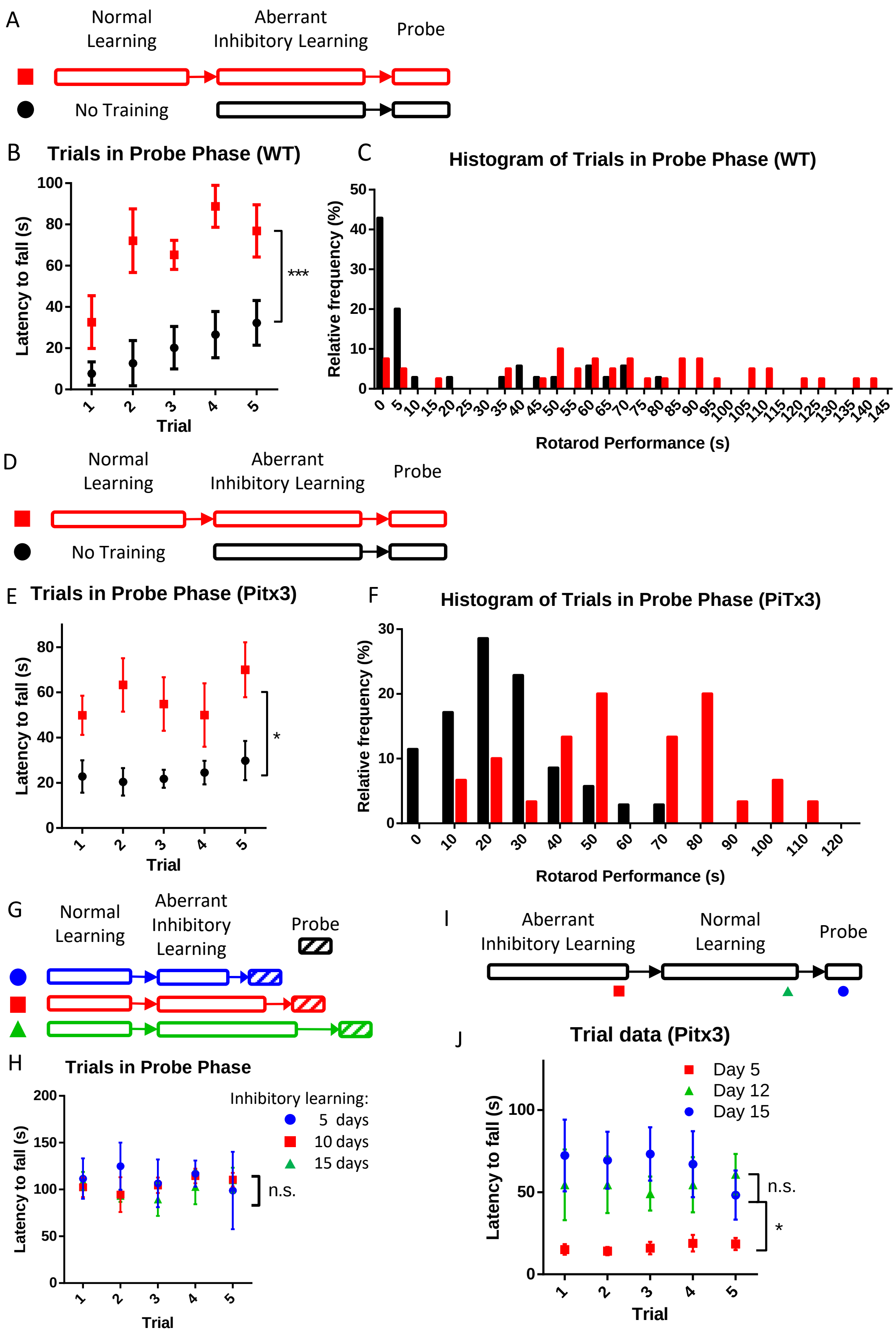

**Fig S1. Individual trial data for different rotarod experiments.**
(A, D, G, I) Behavior paradigms used in each experiment and different phases of normal learning and aberrant inhibitory learning.
(A-C) Preserved normal memory after aberrant inhibitory learning in WT mice.
(B) Individual trial data from figure 1C on day 15, two-way ANOVA, group effect $F (1, 13) = 20.35$, $p = 0.0006$; group x time interaction, $F (4, 52) = 1.239$, $p = 0.3058$.
(C) Histogram of rotarod performance on probe phase.
(D-F) Preserved normal memory after aberrant inhibitory learning in Pitx3 mutant mice.
(E) Individual trial data from day 15 in figure 1D, two-way ANOVA, group effect $F (1, 11) = 8.317$, $p = 0.0149$, group x time interaction, $F (4, 44) = 1.587$, $p = 0.1945$.
(F) Histogram of rotarod performance on probe phase.
(G-H) Preserved normal memory after various days of aberrant inhibitory learning in Pitx3 mutant mice.
(H) Individual trial data from probe phase after various days of aberrant inhibitory learning from figure 1E. Two-way ANOVA between three groups with different length of inhibitory learning, group effect $F (2, 17) = 0.255$, $p = 0.778$; group x time interaction $F (8, 68) =0.372$, $p = 0.932$.
(I-J) Inhibitory memory was reversed by normal learning experience in Pitx3 mutant.
(I) Experimental design with red, green and blue symbols showing different time points for comparison.
(J) Individual trial data on day 5, 12, 15 from figure 1F. Two-way ANOVA with repeated measures between day 5 and day15, day effect $F (1, 7) = 11.80$, $p = 0.022$. Two-way ANOVA with repeated measures between day 12 and day15, day effect $F (1, 7) = 3.650$, $p = 0.195$. P values are adjusted using Bonferroni correction.
All data represents mean ± SEM. *, $p<0.05$; **, $p<0.01$; ***, $p<0.001$. n.s., not significant.

Supplementary Figure 2

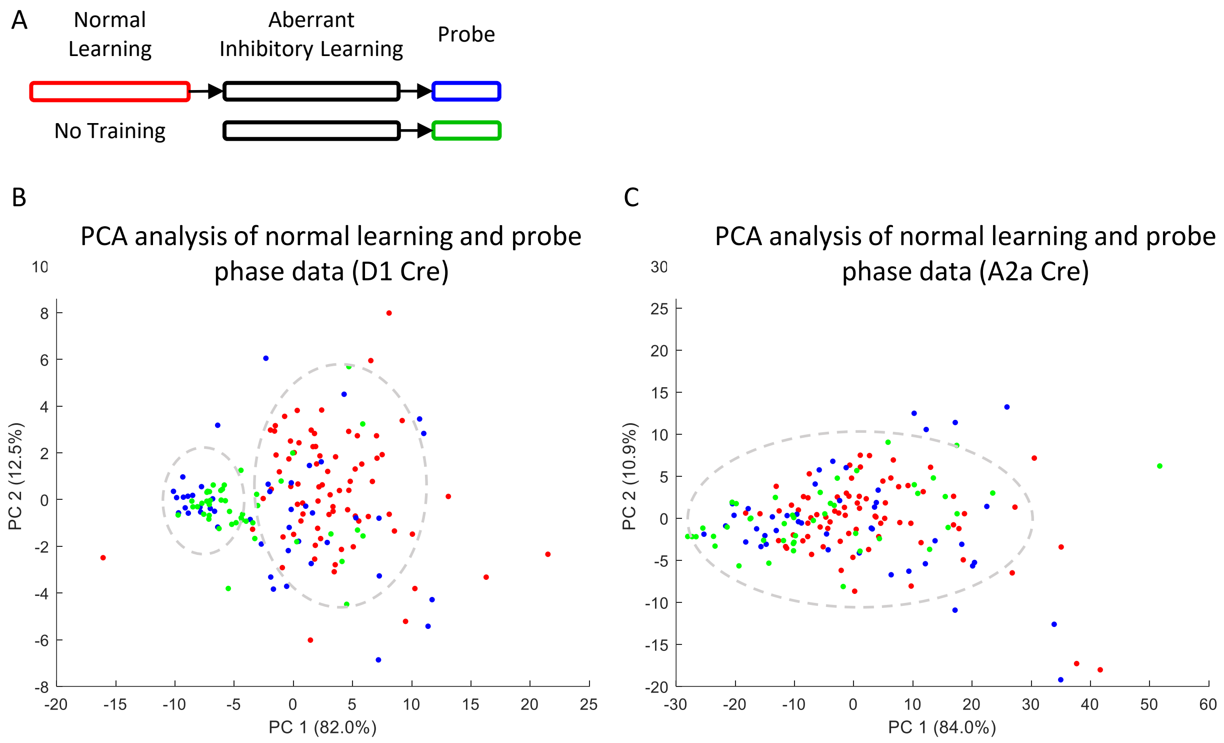


**Figure S2. PCA analysis of fiber photometry signal**

(A) Behavior experiment design and color code for data points from different phases. The red, blue and green section shows where the data were collected in fig S2B-C.

(B) Plotting dSPNs fiber photometry data using principal component 1 and 2 from PCA analysis. Red, normal learning data; blue, probe phase data from the group trained in normal learning phase; green, probe data from the group not trained in normal learning phase.

(C) Plotting iSPNs fiber photometry data using principal component 1 and 2 from PCA analysis. Color codes are the same as figure S2A.

Supplementary Figure 3

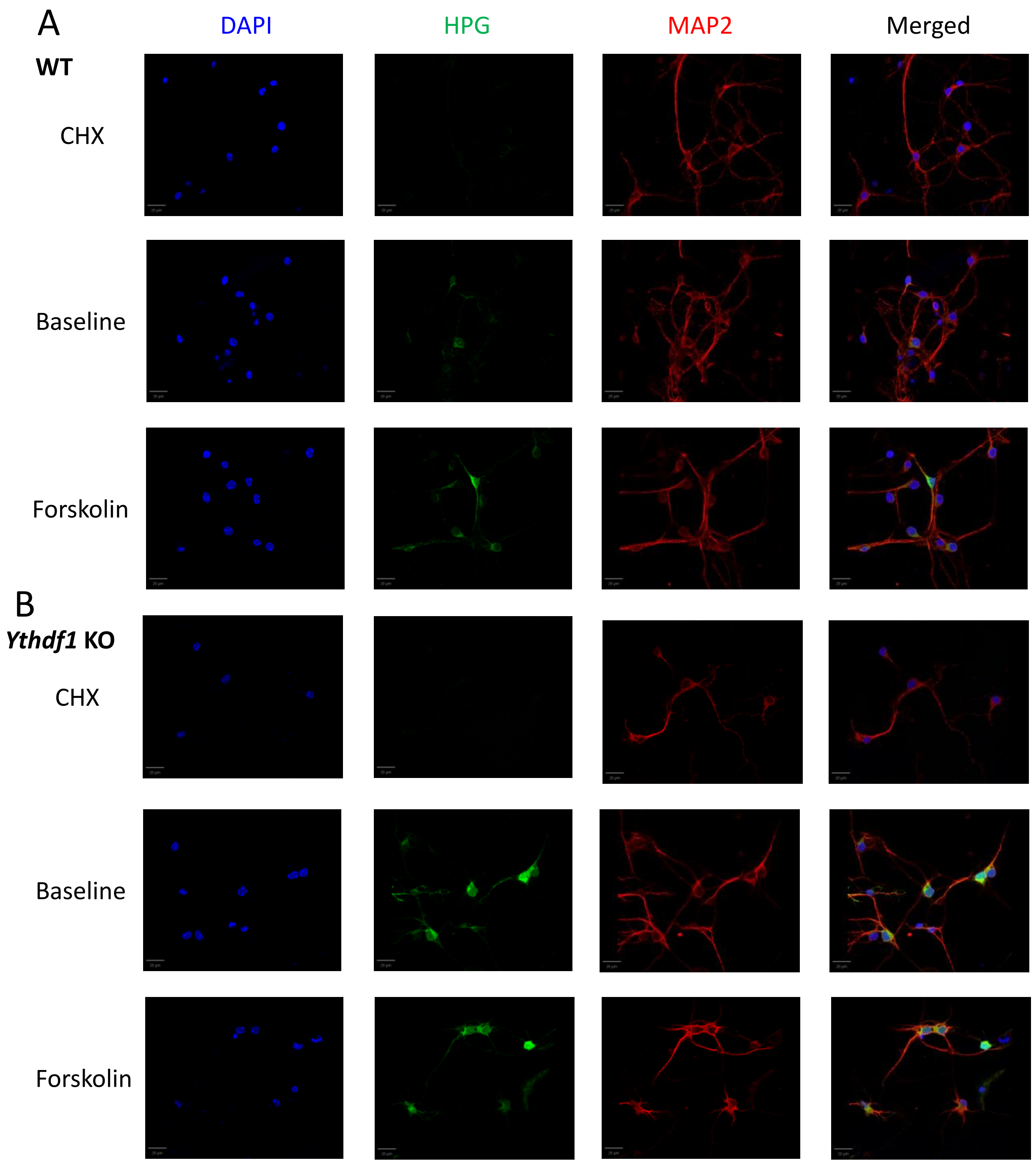

Supplementary Figure 3 (continued)

C

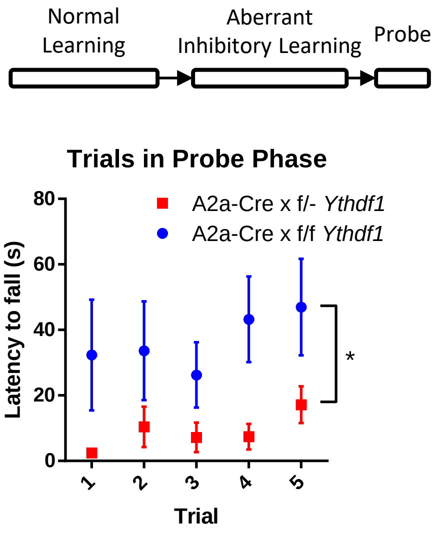


**Figure S3. Measuring new protein synthesis in *Ythdf1* KO using Click chemistry.**

(A) Newly synthesized protein in WT control. Left-right: DAPI, HPG signal, MAP2, merged; top-bottom: CHX treated, baseline and forskolin treated group. Scale bar, 20um.

(B) Newly synthesized protein in *ythdf1* KO. Left-right: DAPI, HPG signal, MAP2, merged; top-bottom: CHX treated, baseline and forskolin treated group. Scale bar, 20um.

(C) Individual trials of rotarod performance from probe phase (day 15 in Figure 4E). Two-way ANOVA, Group effect, $F(1, 9) = 5.764$, $p=0.0398$. Group x time interaction $F(4, 36) = 0.5931$, $p=0.6698$. n(A2a Cre x f/f *Ythdf1*) = 5. n(A2a Cre x f/- *Ythdf1*) = 6.

Supplementary Figure 4

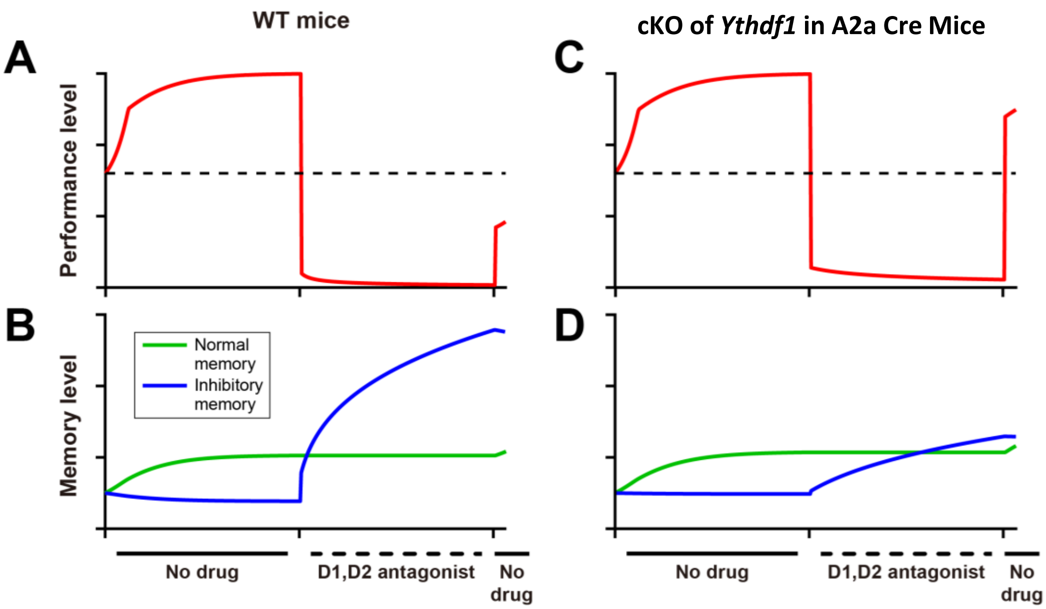


**Figure S4. Computational model describing rotarod motor learning and performance in mice with conditional *Ythdf1* gene deletion in A2a Cre mice.**
(A) Behavior of WT mice predicted by the model in a "normal learning - aberrant inhibitory learning - probe" design. (B) The weights of normal memory and aberrant inhibitory memory in the model. (C – D) Similar as (A – B) but in mice with A2a-Cre mediated conditional *Ythdf1* gene deletion where the inhibitory learning rate becomes 10% of the normal rate.

### Supplemental Information: Details of the basal ganglia computational model

#### The "cortico-basal ganglia-cortical" circuitry

Our basal ganglia model incorporates interactions among the motor cortex and D1, D2 nuclei. In order to ease our analysis, we model the population mean activity of these areas, given by scalar variables $[Cx]$, $[D1]$ and $[D2]$. To be specific, $[Cx]$ represents the mean activity of the specific part (sub-population or sub-dimension) of the motor neuron population that controls the rotarod task. Similarly, $[D1]$, $[D2]$ represent the specific part of dSPN, iSPN neuron population that controls the rotarod task, respectively. Our aim is to recapitulate the latency of the animal's fall during the rotarod task in the experimental data, with the dynamics of $[Cx]$ in the model, which reflects the animal's behavioral performance for the task. Based on the simplified "cortico-basal ganglia-cortical" architecture (Figure 5A), we propose that $[Cx]$ activates $[D1]$ and $[D2]$ with effective connection weights $W_{1C}$ and $W_{2C}$ respectively; $[D1]$ activates $[Cx]$ with effective weight $W_{C1}$, forming a positive feedback loop; while $[D2]$ inactivates $[Cx]$ with effective weight $W_{C2}$, forming a negative feedback loop. $W_{1C}$ and $W_{C1}$ represent the level of normal memory stored in the direct pathway, while $W_{2C}$ and $W_{C2}$ represent the level of inhibitory memory stored in the indirect pathway. Overall, the temporal dynamics of $[Cx]$, $[D1]$ and $[D2]$ are determined by these connection weights with the following set of differential equations:

$$
\begin{aligned}
\frac{d[\mathrm{D}_1]}{dt} &= -[\mathrm{D}_1] + f(W_{1\mathrm{C}} \cdot [\mathrm{Cx}] \cdot G_1^A), \\
\frac{d[\mathrm{D}_2]}{dt} &= -[\mathrm{D}_2] + f(W_{2\mathrm{C}} \cdot [\mathrm{Cx}] \cdot G_2^A), \\
\frac{d[\mathrm{Cx}]}{dt} &= -[\mathrm{Cx}] + f(W_{\mathrm{C}1} \cdot [\mathrm{D}_1] - W_{\mathrm{C}2} \cdot [\mathrm{D}_2] + I_{\mathrm{Motion}}).
\end{aligned} \tag{1}
$$

Here $I_{\mathrm{Motion}} = 1$ represents a constant external drive to the motor cortex to start the motor programs associated with the rotarod task. The coefficents $G_1^A$ and $G_2^A$ represent the dopamine dependent gain for how $[Cx]$ activity drives $[D1]$ and $[D2]$ activity. Since dopamine activates D1 pathway and inhibits D2 pathway, we set $G_1^A = 2$; $G_2^A = 0.5$ for the wild type animal with normal dopamine level, or Pitx3 mutant mice but treated with L-DOPA. While when antagonists of D1, D2 receptors are applied in the wild type animal, we set $G_1^A = 0.1$; $G_2^A = 10$ to simulate the effect of D1 inhibition and D2 disinhibition by the antagonist. Finally, the function $f(x)$ represents the activation function:

$$
f(x) = \begin{cases} 0 & (x <= 0) \\ x & (0 = x <= C_{\max}) \\ C_{\max} & (x > C_{\max}) \end{cases} \tag{2}
$$

The function $f(x)$ sets up lower and upper bounds for the total external input to $[Cx]$, $[D1]$ and $[D2]$, with the threshold $C_{\max} = 3$.

Finally, since the temporal dynamics of neuronal activities in the Eq. (1) are much faster than long-term plasticity and learning process, we only consider the steady state solutions of the Eq. (1) (where the left hand sides become 0).

#### Long-term plasticity rules of the normal memory and inhibitory memory

We propose that the dopamine regulated long-term plasticity processes, i.e. the dynamical rules governing the evolution of $W_{1C}$, $W_{C1}$, $W_{2C}$ and $W_{C2}$, are based on the "cAMP-protein synthesis-memory consolidation" process (Figure 5B). For the weights of the direct pathway ($W_{1C}$, $W_{C1}$) that interconnects [Cx] and [D$_1$], we propose the following learning rule:

$$\frac{dW_{\mathrm{1C}}}{dt} = \frac{dW_{\mathrm{C1}}}{dt} = \eta \cdot G^L \cdot (\mathrm{C_{max}} - [\mathrm{Cx}]) \cdot ([\mathrm{Cx}]. \cdot [\mathrm{D_1}]). \tag{3}$$

We note that the dynamics of $W_{1C}$ and $W_{C1}$ are symmetric, and the initial states are $W_{1C}(0) = W_{C1}(0) = 0.5$, implying that $W_{1C}(t) = W_{C1}(t)$ for all $t$. Here $G^L = 1$ or 0 is a binary gating variable of dopamine regulation on long-term plasticity: $G^L = 1$ when dopamine level is normal, such as for wild type mice or Pitx3 mutant mice but treated with L-DOPA. By contrast $G^L = 0$ when dopamine regulation is absent, such as when the antagonist is applied to the wild type mice, or L-DOPA is absent for Pitx3 mutant mice. This models that a lack of dopamine blocks the cAMP dependent protein synthesis and the following normal learning process (Figure 5B). The dynamics of $W_{\mathrm{C1}}$ is given by the product of the prediction error term $\mathrm{C_{max}} - [\mathrm{Cx}]$, and the Hebbian learning term $[\mathrm{Cx}] \cdot [\mathrm{D_1}]$, since we propose that long-term plasticity in the D1 pathway occurs only when there are reward prediction errors[1]. Given that animals are expected to fully acquire the running skill to avoid falling when performing the rotarod task, we propose that the reward prediction error is proportional to $\mathrm{C_{max}} - [\mathrm{Cx}]$, i.e. a linear decay with the performance level that stay maximum without any learning, and become zero for optimized performance. Thus, an increase in the motor activity and behavioral performance ([Cx]) due to the dopamine-driven LTP ($G^L = 1$) will decrease the reward prediction error and slow down the plasticity, until the optimal performance is achieved ($[\mathrm{Cx}] = \mathrm{C_{max}}$) and reward prediction error becomes 0. We set $\mathrm{C_{max}} = 3$, being the same as the threshold in Eq. (2), and [Cx] and [D$_1$] are given by the steady state solution of the fast firing rates in Eq. (1), as we mentioned above. This requires the learning rate $\eta$ to be small; we set $\eta = 2 \times 10^{-3}$.

For the indirect pathway ($W_{2C}$, $W_{C2}$) that interconnects [Cx] and [D$_2$], we propose the following learning rule:

$$\frac{dW_{\mathrm{2C}}}{dt} = \frac{dW_{\mathrm{C2}}}{dt} = \eta \cdot \begin{cases} -(\mathrm{C_{max}} - [\mathrm{Cx}]) \cdot ([\mathrm{Cx}] \cdot [\mathrm{D_2}]) & (G^L = 1) \\ G_2^A \cdot ([\mathrm{Cx}] \cdot [\mathrm{D_2}]) & (G^L = 0). \end{cases} \tag{4}$$

Here $G^L$ is the same as in the Eq. (3). When dopamine regulation exists ($G^L = 1$), the learning rule would be similar to that in the Eq. (3) but reversed (the $-1$ coefficient), since dopamine drives LTD (rather than LTP) of the indirect pathway. By contrast, when dopamine is absent ($G^L = 0$), the cAMP in D2 neurons will no longer be suppressed, so that the learning of the indirect pathway will be disinhibited, i.e. the inhibitory memory level will increase. Since dopamine is absent, the learning will no longer be modulated by reward prediction error, so the learning rule will become a simple Hebbian form ($[\mathrm{Cx}] \cdot [\mathrm{D_2}]$). The term $G_2^A$ is the same as in the Eq. (1), which equals to 10 when dopamine is absent.

All learning simulations have 100 iteration steps for each learning phase, i.e. 300 steps totally for groups with "normal - aberrant inhibitory - probe" schedule, and 200 steps for groups with "aberrant inhibitory - probe" schedule (red and black curves in Figure 5D and 5H).

##### Model performance for the "normal learning - aberrant inhibitory learning - probe" schedule in the wild-type mice

Figure 5D, 5H and 5L show the simulation results of [Cx] over the learning process, and Figure 5E-5F, 5I-5J and 5M show the dynamics of $W_{1C}$, $W_{C1}$ (i.e. normal memory in the direct pathway) and $W_{2C}$, $W_{C2}$ (i.e. aberrant inhibitory memory in the indirect pathway). For normal learning in the WT mice without antagonist drugs (Figure 5D-5E, red group), dopamine activates LTP of the direct pathway and LTD of the indirect pathway (Figure 5E, increase and decrease for the green and blue curves), until the performance [Cx] approaches the optimal value $C_{max}$ (Figure 5D, saturation of the red curve) so that the reward prediction error becomes 0 and learning stops (Figure 5E, saturation of green and blue curves). When antagonists are applied at the beginning of the aberrant inhibitory learning phase, the gain values $G_1^A$, $G_2^A$ that dopamine modulate the [$D_1$] and [$D_2$] activity are changed (from $G_1^A = 2$; $G_2^A = 0.5$ to $G_1^A = 0.1$; $G_2^A = 10$), leading to an immediate drop of the steady state solution of [Cx] (Figure 5D, drop in the red curve). Then, the gating variable of dopamine regulation on long-term plasticity, $G^L$, is switched from 1 to 0, so that as time goes during the aberrant inhibitory learning phase, the long-term plasticity in the direct pathway stops (Figure 5E, green curve keep steady), but a strong LTP happens for the indirect pathway (Figure 5E, increasing blue curve). When the system skips the normal learning during the first phase (Figure 5D, 5F, black group), the LTP of normal memory during the normal learning phase will not happen (Figure 5E-5F, compare the levels of the green curves during the beginning of aberrant inhibitory learning phase ), so that a much weaker $W_{1C}$, $W_{C1}$ leads to a dramatic difference of task performance [Cx] when probe phase begins (Figure 5D, compare red and black at probe phase).

##### Model performance for the "normal learning - aberrant inhibitory learning - probe" schedule in the Pitx3 mutant mice: Adaptation of dopamine receptors

For Pitx3 mutant mice, the learning dynamics when external L-DOPA applied is similar to that of the normal WT mice (where $G^L = 1$), and learning without L-DOPA drug is similar to that of WT mice with antagonists (where $G^L = 0$). As a result, the dynamics of memory change would be similar for WT mice and Pitx3 mutant mice (Figure 5E-5F vs 5I-5J). Also, the gain values $G_1^A$ and $G_2^A$ that regulate [$D_1$] and [$D_2$] activities are also the same for external L-DOPA applied in mutant mice and normal WT mice ($G_1^A = 2$; $G_2^A = 0.5$). However, we propose that when L-DOPA is absent, these gain values are different from the situation when antagonists are applied in WT mice. Since Pitx3 mutant mice lacks internal expression of dopamine at early development, we assume that D1 and D2 receptors would become adapted to the environment of low level of dopamine. As a result, D1 receptors become ultra-sensitive to dopamine, so that they will keep slightly activated without external L-DOPA; and D2 receptors become insensitive to the lack of dopamine, so that they will somewhat keep suppressed without external L-DOPA. Thus, we set $G_1^A = 1.8$; $G_2^A = 0.8$ for Pitx3 mutant mice without external L-DOPA, which is similar to the situation when external L-DOPA applied but much different from when antagonist applied in WT mice. As a result, the performance decays smoothly during the aberrant inhibitory learning phase in Pitx3 mutant mice (Figure 5H), rather than an instant drop in WT mice (Figure 5D). For a similar reason, in the 'aberrant inhibitory learning-normal learning-probe' schedule (Figure 5L-5M), the performance would persist right after L-DOPA removal at the beginning of the probe phase (Figure 5L).

#### Learning in the mice with A2a-Cre conditional KO of *Ythdf1*

For the performance of A2a-Cre x floxed-*Ythdf1* mice (Figure 4 and Supp Figure 4), we proposed that the YTHDF1 knock-out in the D2 pathway would limit the proteins synthesis process that are necessary for the consolidation of aberrant inhibitory memory. Thus, we assume that the inhibitory learning would be slower in the A2a-Cre x *Ythdf1* mice than that in WT animal. As a result, the normal-aberrant inhibitory memory balance would be more biased to the normal memory side, leading to a higher performance level right after antagonist removal than that of the control group in the model (Supp Figure 4A-4B vs 4C-4D).

#### Difference of D1, D2 unit activities recorded in fiberphotometry experiments and in the model

For the fiberphotometry recording of population mean activity of dSPN (D1) and iSPN (D2), we note that the activity of dSPN and iSPN population is anti-correlated to the performance during the normal learning process (Figure 2E, 2G), which, is not the case for our model. During the normal learning process of WT animals, the level of performance [Cx] and normal memory $W_{1C}$, $W_{C1}$ will increase, and the level of inhibitory memory $W_{2C}$, $W_{C2}$ will decrease. Overall, this leads to an increase for both [$D_1$] and [$D_2$] activity in the steady state solutions of the Eq. (1). However, the fiberphotometry imaging records the mean activity of a large dSPN, iSPN population, while the variable [$D_1$] and [$D_2$] in the model represents the certain sub-population / sub-dimension of dSPN and iSPN that specifically control the rotarod task. The activation of dSPN and iSPN sub-population activities for the rotarod task leads to a selection of the rotarod motion program over the other programs in the motor cortex, possibly through mutual inhibition among different sub-populations of motor cortex, dSPN and iSPN for different motion programs. Thereby, we proposed that the remaining sub-populations that are relevant to all the other programs will be largely suppressed, leading to a overall decrease of the mean activity of the whole dSPN, iSPN population, as in the experiment (Figure 2E, 2G). While the significant changes of dSPN, iSPN activity during the learning process still provide strong neuronal-level evidence that the long-term plasticity related to dSPN and iSPN does happen during the animal training.
